# Scalable expansion of human iNKT cells: single-cell profiling and in vivo control of GvHD with preserved GvL activity

**DOI:** 10.64898/2026.09.11.750594

**Authors:** Jordan Brouard, Cristina Caraiman, Ghislain Fievet, Lucile Monchablon, Tereza Coman, Sebastien Hergalant, Simona Pagliuca, David Moulin, Marie Thérèse Rubio

## Abstract

Invariant natural killer T (iNKT) cells can limit graft-versus-host disease (GVHD) after hematopoietic stem cell transplantation (HSCT), but their scarcity in peripheral blood limits therapeutic development. Current clinical-grade human iNKT expansion protocols mainly rely on IL-2, require prior iNKT-cell sorting, last 6–8 weeks, and predominantly expand CD4+ iNKT cells, whereas human CD4^−^ iNKT cells are more strongly associated with GVHD control in patients and uniquely regulate antigen-presenting cells and T-cell activation. We developed a scalable culture system to preferentially expand human CD4^−^ iNKT cells directly from total peripheral blood mononuclear cells (PBMCs) using alpha-galactosylceramide (α-GalCer) and optimized cytokine conditions. IL-15 was the most effective cytokine. The optimized 14-day protocol generated a mean of 3.8×10^7^ iNKT cells from 2×10^7^ PBMCs, including 74% CD4^−^ iNKT cells. Single-cell transcriptomic profiling identified eight major iNKT subsets, differentiation trajectories during expansion, and distinct IL-2- versus IL-15-associated transcriptional programs. IL-15-expanded iNKT cells induced apoptosis of monocyte-derived dendritic and leukemic cells in vitro, controlled xeno-GVHD, and preserved graft-versus-leukemia (GVL) activity in preclinical mouse models. This platform enables reproducible production of human CD4^−^ iNKT cells at clinically relevant scale and position IL-15–expanded iNKT cells as a compelling immunotherapy candidate for allo-HSCT.

## Introduction

Invariant Natural Killer T cells (iNKT) are a rare highly conserved subset of T lymphocytes with pivotal immunomodulatory properties. They express an semi-invariant T-cell receptor (TCR) composed of Vα24-Jα18 paired with Vβ11 in humans, which specifically recognizes glycolipids presented by CD1d molecules ^1,2^. In humans, numerous studies have highlighted the importance of iNKT cell levels, functionality and subsets in tumor immunosurveillance, autoimmune diseases prevention and clearance of infectious pathogens ^3–6,2^.

Allogeneic hematopoietic stem cell transplantation (allo-HSCT) stands as a complex cellular therapy hinging on the anti-tumoral effect elicited by donor immune cells. The benefice of the graft versus tumor (GVT) effect is counterbalanced by the onset of graft-versus-host disease (GvHD) causing critical organ toxicity^7^. Despite extensive pharmaceutical endeavors, acute GvHD afflicts 30–50% of patients constituting a major cause of morbidity and mortality^8^. In parallel, disease relapse remains the foremost mortality cause after allo-HSCT ^9^.

Our research along others, has demonstrated that higher levels of human iNKT cells, particularly the CD4^−^ subset correlate with diminished GVHD incidence while simultaneously preserving GVT efficacy ^10–12^. These clinical observations have been supported by both *in vitro* and *in vivo* studies using GVHD models, confirming the unique protective role of human CD4^−^ iNKT cells ^13–15^.

Given their broad immunoregulatory functions, iNKT cells are promising candidates for cellular immunotherapy, particularly in the allo-HSCT setting. However, their very low frequency in peripheral blood (∼0.001–1%) and marked inter-donor variability make clinical-scale expansion challenging, especially for autologous applications. Current GMP-grade expansion methods typically require initial iNKT cell sorting from PBMCs, feeder cell support, 6–8 weeks of culture, and preferentially expand CD4+ iNKT cells^16–20^. Alternative approaches rely on genetically engineered hematopoietic or pluripotent stem cells ^15,21,22^.

Here, we describe a simpler, feeder-free, and initially sorting-free ex vivo expansion protocol that preferentially expands the CD4^−^ iNKT cell fraction. Using single-cell RNA sequencing, we characterized the final product and provided the first comparison of the iNKT subtypes generated under ex vivo expansion conditions. We also present the first preclinical evidence that IL-15-expanded human iNKT cells can control GvHD while preserving the GvL effect in a xeno-GVHD model. Together, these findings support a promising and scalable strategy for future cellular immunotherapy applications.

## Results

### Optimized culture conditions promoting CD4− iNKT cell expansion

Invariant NKT lymphocytes can be expanded in vitro by culturing PBMC in RPMI medium supplemented with 10% fetal calf serum (FCS), alpha-galactosylceramide, a synthetic glycolipid that activates iNKT cells through their invariant T-cell receptor, and interleukin-2 (IL-2). Although this protocol supports iNKT cell expansion for functional studies, it does not preferentially expand CD4^−^ iNKT cells or yield sufficient cell numbers for therapeutic use^13^.

We first sought to optimize the expansion of this subset from healthy donor PBMCs under small-scale culture conditions (2 mL) by comparing IL-2, IL-4, IL-7, and IL-15, used alone or in combination in addition to α-GalCer, over 14 days, with or without a medium change on day 7 (**Figure 1A**). We assessed the effect of each cytokine on the iNKT-cell expansion factor, defined as the ratio of iNKT cells recovered on day 14 to those present on day 0. IL-15 markedly increased the expansion of CD4^−^ iNKT cells (mean [SD], 691.5 [1187]) compared with IL-2 (78.65 [142.6], p=0.0045), IL-7 (24.17 [64.77], p=0.0001), and IL-4 (3.012 [4.012], p<0.0001) (**Figure 1C**). The beneficial effect of IL-15 was also observed in CD4^+^ iNKT cells, with a significantly greater expansion factor than with IL-2 (p=0.0107), IL-7 (p=0.0035), or IL-4 (p=0.0002) (**Figure 1C**). In contrast, combining IL-15 with IL-2, IL-4, and/or IL-7 did not significantly improve the expansion factor of CD4^−^ iNKT cells compared with IL-15 alone (**Figure S1A**). Under the IL-15-alone condition, the median proportion of double-negative CD4/CD8 (DN) iNKT cells reached 54.87% [SD 22.19], significantly higher than with IL-7 (34.27% [SD 21.34], p=0.0029) or IL-4 (32.02% [SD 13.82], p=0.0171) (**Figure 1D**). IL-15 also yielded the highest number of CD4^−^ iNKT cells, compared with IL-2 (p=0.0002) (**Figure 1E**).

**Figure 1:**
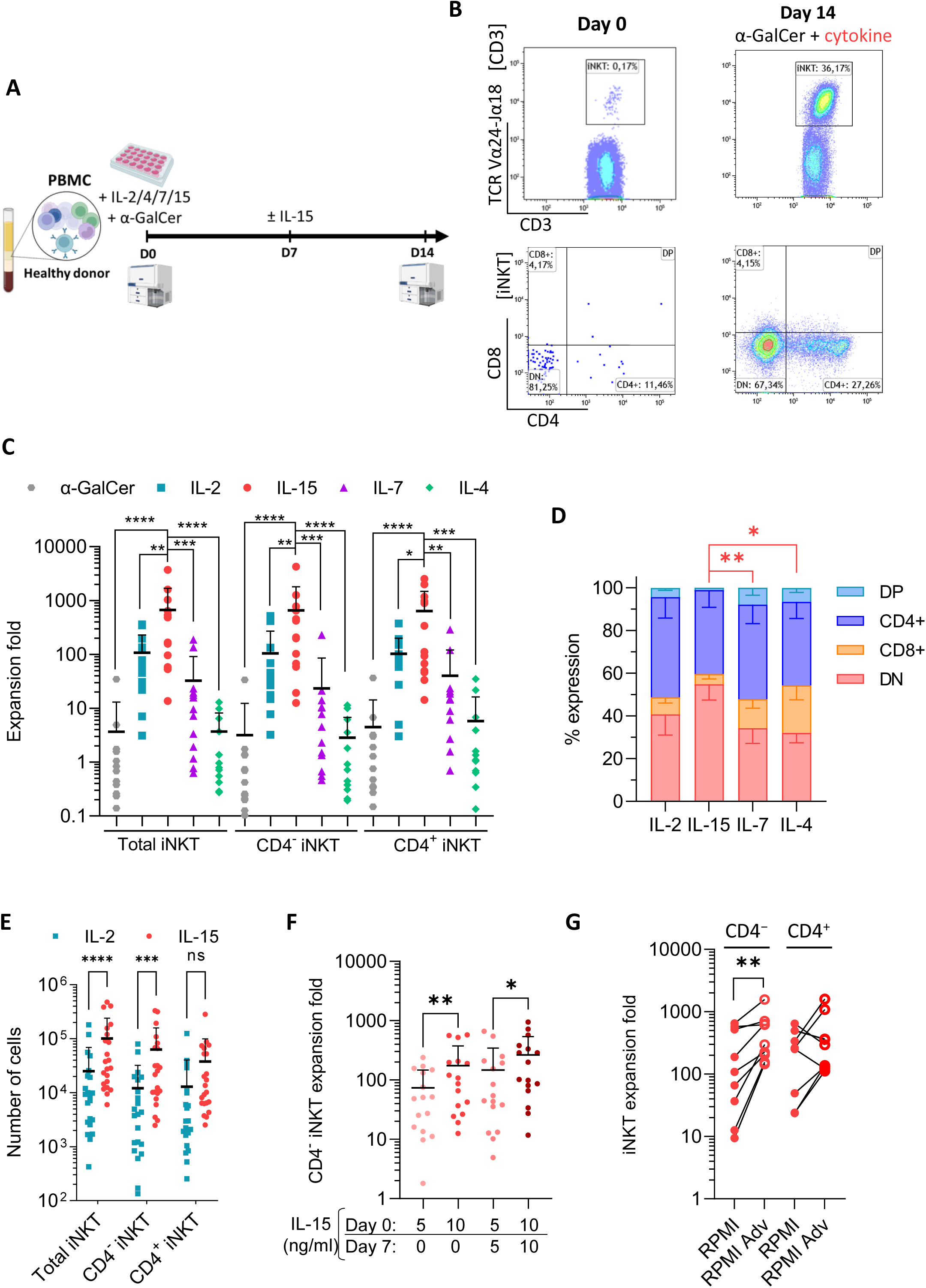
IL-15 preferentially drives the expansion of human CD4− iNKT cells. (A) Schematic overview of the human iNKT-cell expansion protocol. PBMCs were cultured from D0 with α-GalCer and the indicated cytokines for 14 days. In selected experiments, half of the medium was replaced on D7 and IL-15 was added. (B) Representative flow cytometry plots from one donor showing the percentage of iNKT cells (6B11+) among CD3+ cells at D0 in PBMCs and after expansion without prior isolation in the presence of α-GalCer and IL-2 (top), together with the corresponding CD4/CD8 phenotype (bottom). (C) Expansion fold from D0 to D14 of total, CD4−, and CD4+ iNKT cells according to the cytokines used (n=13). Data are shown as mean + SD; comparisons were performed using ANOVA with Friedman’s test. (D) Distribution of iNKT cells according to CD4, CD8, double-negative (DN), or double-positive (DP) phenotype under each cytokine condition (n=9). Data are shown as mean - SEM; comparisons were performed using two-way ANOVA. (E) Number of totals, CD4−, and CD4+ iNKT cells after expansion with IL-2 (blue squares) or IL-15 (red circles) (n=22). Data are shown as mean + SD; comparisons were performed using two-way ANOVA. (F) Expansion fold of CD4− iNKT cells according to IL-15 concentration (5 or 10 ng/mL), with or without additional cytokine supplementation on D7 (n=16). Data are shown as mean + SD; comparisons were performed using two-way ANOVA. (H) Expansion fold of CD4− and CD4+ iNKT cells according to the culture medium used, conventional RPMI or RPMI Advanced (Adv) (n=9). Comparisons were performed using the Wilcoxon test.

Because IL-15 showed the strongest effect on CD4^−^ iNKT-cell expansion, we next evaluated different doses and schedules of IL-15 supplementation. A single addition of IL-15 at 10 ng/mL on day 0 produced greater expansion than 5 ng/mL on day 0 (p=0.0061) and performed similarly to 5 ng/mL given on days 0 and 7 (p=0.7920) or 10 ng/mL given on days 0 and 7 (p=0.7920) (**Figure 1F**). Higher doses of 20 or 50 ng/mL did not provide any further benefit (not shown).

Lastly, we observed that RPMI Advanced (RPMI Adv) also significantly improved CD4^−^ iNKT-cell expansion compared with conventional RPMI, leading to more than 100-fold expansion of CD4^−^ iNKT cells in all samples (min-max: 9.4-650.4 for RPMI and 145.3-1579 for RPMI Adv, p=0,0039) (**Figure 1G**). Overall, the optimal conditions for preferential CD4^−^ iNKT cell expansion consisted of culturing PBMCs for 14 days in RPMI Advanced supplemented on day 0 with 5% SVF, IL-15 (10 ng/mL), and α-GalCer (100 ng/mL).

### High scale in vitro expansion can achieve sufficient iNKT absolute numbers for therapeutic use

After small-scale culture experiments, we assessed intermediate-scale expansion using either culture bags (5×10^7^ PBMC in 100 mL) or G-Rex bioreactors (2×10^7^ PBMC in 100 mL) under the optimized culture conditions defined above (**Figure 2A**). iNKT cells expanded in both systems, but the proportion of CD4^−^ iNKT cells was significantly higher during the expansion in G-Rex bioreactors, rising from 55.16% (SD: 20.16) at day 0 to 67.27% (SD: 23.86) at day 14 (p = 0.0444). By contrast, the proportions of CD4^−^ and CD4^+^ iNKT cells remained unchanged in culture bags (**Figure 2B**). G-Rex bioreactors also yielded significantly greater expansion than culture bags, for CD4^−^ iNKT cells (mean expansion factor: 2331.4 [SD 2819.8] vs 273.7 [SD 358.7], respectively; p= 0.0088) but also for CD4^+^ iNKT cells (p= 0.0088) (**Figure 2C**). To define the optimal culture duration, we monitored iNKT-cells cultured for up to 21 days, with or without the addition of PBMCs, α-GalCer, and IL-15 on day 14. In both conditions, iNKT cell numbers peaked at day 14 and then declined. This kinetic profile was consistent with the crossover in glucose consumption and lactate production observed at day 14, supporting a 14-day culture period as optimal (**Suppl Figures 1C and 1D**). The optimized 14-day G-Rex protocol with IL-15 generated a mean of 4.1×10^7^ total iNKT cells (SD: 5.1×10^7^), including 2.84×10^7^ CD4^−^ iNKT cells (SD: 3.8×10^7^), from 2×10^7^ PBMCs. In comparison, IL-2 produced fewer CD4^−^ iNKT cells (mean, 2.12×10^7^; SD: 2.5×10^7^; p=0.0083), while CD4^+^ iNKT cell numbers did not differ between conditions (p=0.9972) (**Figure 2D**).

**Figure 2:**
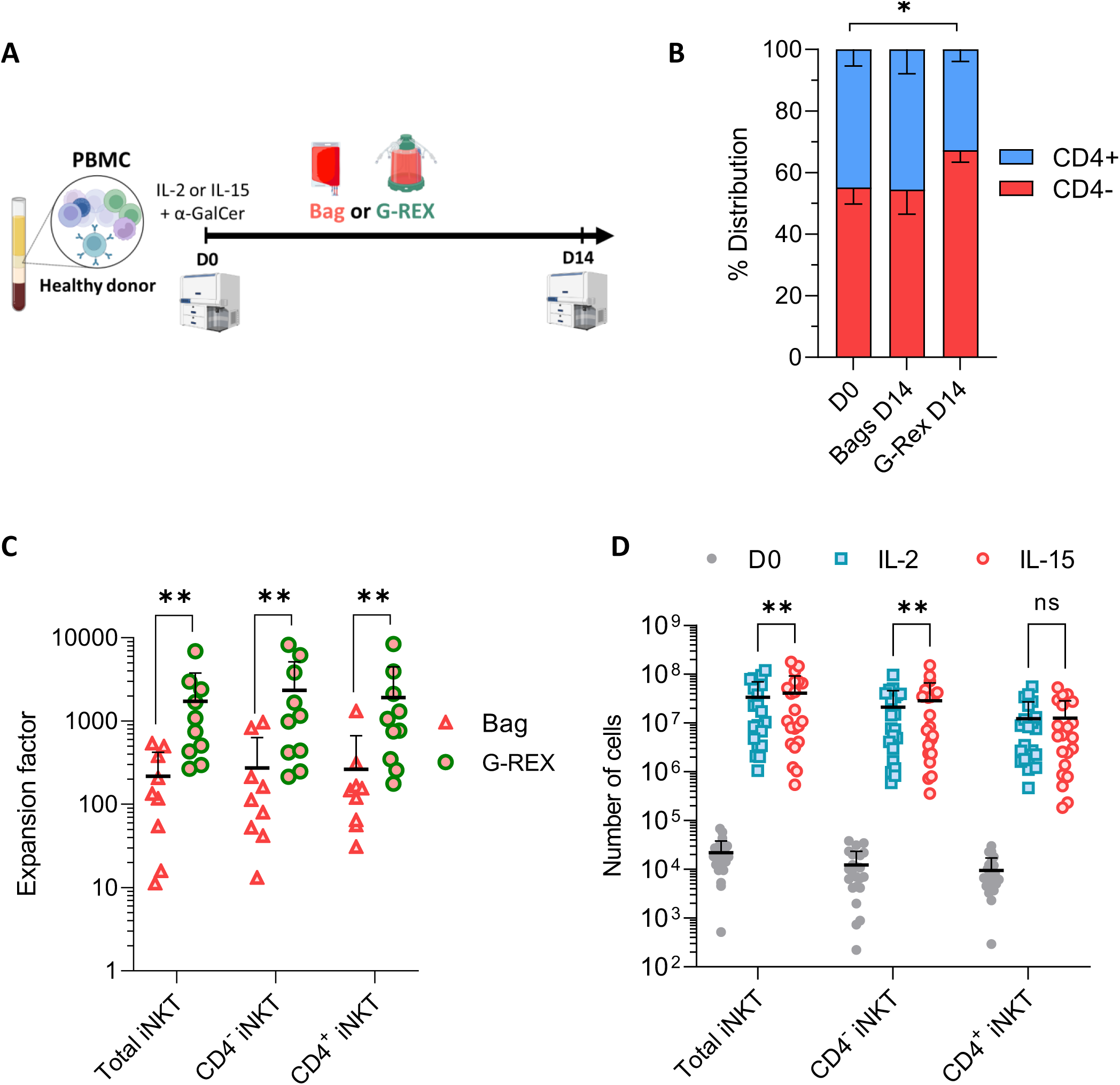
G-Rex bioreactors support large-scale expansion of CD4− iNKT cells. (A) Schematic overview of the large-scale iNKT-cell expansion protocol. PBMCs were cultured from D0 with α-GalCer and IL-15 in culture bags or G-Rex bioreactors for 14 days without intervention. (B) Expansion fold of total, CD4−, and CD4+ iNKT cells according to the culture vessel used, bag or G-Rex (n=9). Data are shown as mean ± SD; comparisons were performed using the Mann-Whitney test. (C) Proportions of CD4− (red) and CD4+ (blue) iNKT cells before expansion (D0, n=14) and after expansion in culture bags (D14 Bag, n=9) or G-Rex bioreactors (D14 G-Rex, n=14). Data are shown as mean ± SEM; comparisons were performed using a paired t-test. (D) Number of total, CD4−, and CD4+ iNKT cells before expansion (D0, grey) and after expansion with IL-15 (red circles) or IL-2 (blue squares) (n=19). Data are shown as mean ± SD; comparisons were performed using paired two-way ANOVA.

### Single-cell characterization of iNKT cells before and after expansion

We next performed single-cell RNA sequencing on baseline iNKT cells (day 0, n=4) and on cells expanded for 14 days in G-Rex with either IL-15 (n=3) or IL-2 (n=3). After immunomagnetic enrichment, cells were stained with an AbSeq panel (CD3, iTCR, CD4, CD8, CD161, CD56) for simultaneous surface protein and transcriptome profiling, then captured using the BD Rhapsody platform (**Suppl Figure 2**). Integration of day 0 and day 14 samples from both cytokine conditions identified eight iNKT clusters and three cycling clusters based on UMAP clustering, AbSeq profiles, and the top five expressed genes in each cluster (**Figure 3A-C**).

**Figure 3:**
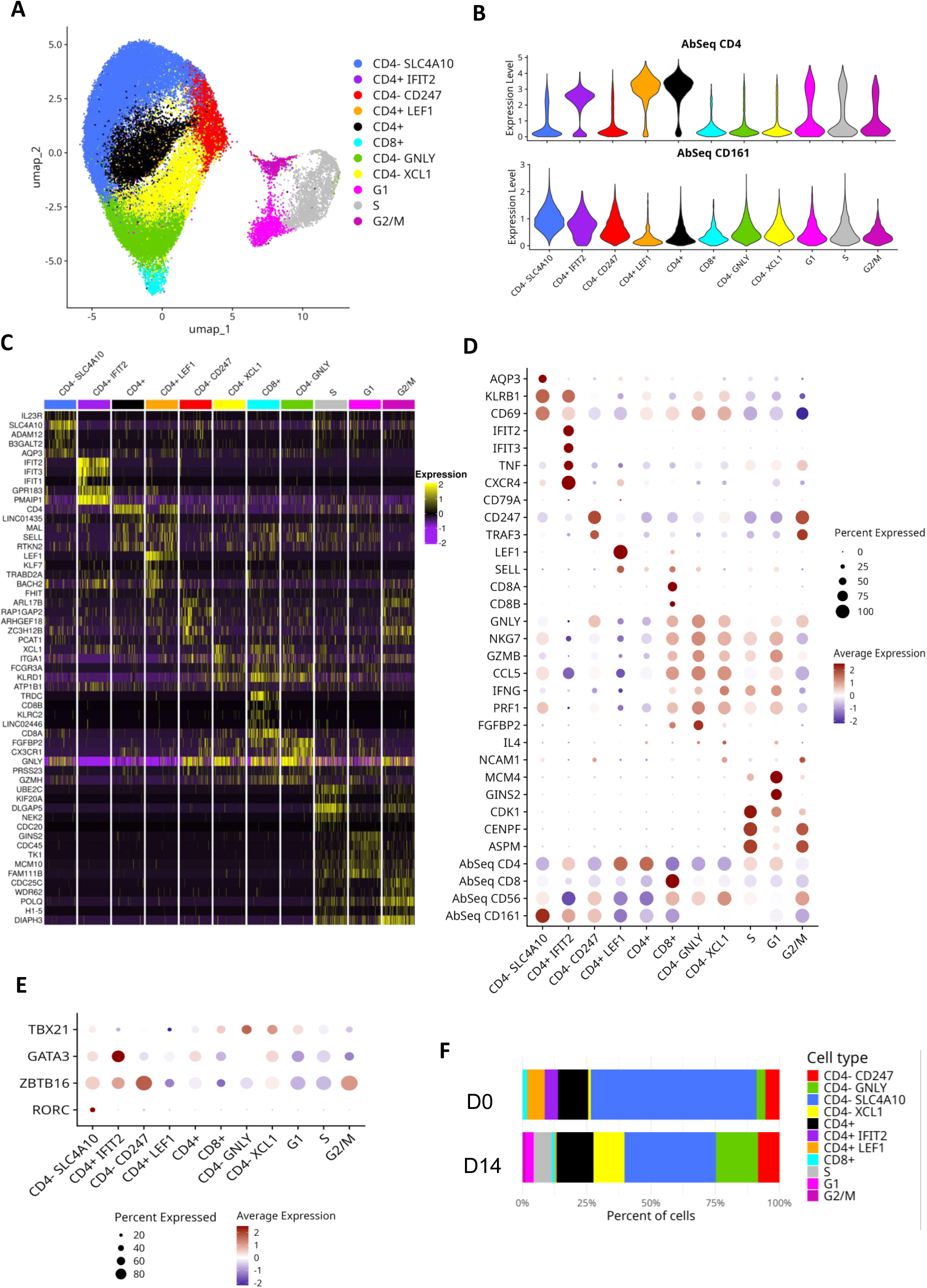
Single-cell RNA-seq reveals marked transcriptomic diversity in human iNKT cells before and after expansion. Sorted human iNKT cells before and after expansion with IL-15 or IL-2 were analyzed by single-cell RNA-sequencing. (A) UMAP of the combined transcriptomic datasets from D0 (n=4), D14 IL-15 (n=3), and D14 IL-2 (n=3), identifying 11 distinct clusters. (B) Violin plots showing CD4 and CD161 AbSeq expression across clusters. (C) Heatmap of the five most highly expressed genes in each cluster. (D) Dot plot showing expression of the indicated genes across clusters. (E) Dot plot showing the expression of lineage-associated genes involved in immune differentiation, including TBX21 (T-bet), GATA3, ZBTB16 (PLZF), and RORC (RORγt), across clusters. (F) Bar plots showing the distribution of iNKT-cell clusters before (D0) and after expansion (D14).

The most abundant population, designated ‘CD4− SLC4A10’, is defined by high *SLC4A10* expression. *SLC4A10* encodes NCBE, a bicarbonate transporter involved in intracellular pH regulation that has previously been described in neurons^23^ and mucosal-associated invariant T (MAIT) cells^24,25^. This population expresses activation markers such as CD69, maturation markers such as KLRB1 (CD161), and cytotoxic markers including *NKG7* and *PRF1*, consistent with an NK-like phenotype (**Figure 3D**). It shows a predominantly Th2 profile, based on *GATA3* and *ZBTB16* (PLZF) expression, but also contains most of the Th17-like cells, identified by *RORC* (RORγ) and *AQP3*, which did not form a distinct cluster (**Figure 3E**).

Among the 4 other CD4 negative/low populations, the ‘CD4^−^ CD247’ cluster shows high expression of CD247 (CD3ζ), a key component of the CD3-TCR complex and a Th2-like iNKT profile (GATA3+, ZBTB16++) (**Figure 3 D-E**)

In contrast, the three other CD4 negative populations: ‘CD8’, ‘CD4^−^ GNLY’, and ‘CD4^−^ XCL1’ display high expression of multiple cytotoxic molecules, including *GNLY* (granulysin), *NKG7*, *GZMB* (granzyme B), and *PRF1* (perforin) and a Th1 profile with high *TBX21* (T-bet) expression (**Figure 3D-E**). These clusters were distinguished by selective CD8A and CD8B expression in the ‘CD8’ cluster, higher FGFBP2 expression in ‘CD4^−^ GNLY’ and higher CD56 expression in the ‘CD4^−^ XCL1’ cluster (**Figure 3D**). Despite this mature phenotype, they show low *KLRB1* transcript levels and weak surface expression of CD161 (**Figure 3B-D**).

Among CD4^+^ iNKT cells, three distinct clusters were identified (**Figures 3B-E**). The ‘CD4^+^ LEF1’ cluster lacks expression of the maturation marker *KLRB1* and is characterized by high *LEF1* and *SELL* (CD62L) expression, without expression of the transcription factors associated with Th1, Th2, or Th17 programs (**Figure 3D-E**). By contrast, the ‘CD4^+^ IFIT2’ cluster displays a mature phenotype (*KLRB1*+) and high expression of genes involved in type I interferon signaling (*IFIT2*, *IFIT3*) and chemokine pathways (*CXCR4*, *CCL5*) (**Figure 3D**). The third cluster ‘CD4^+^’, expressed CD4 together with intermediate *KLRB1* levels (**Figure 3D**). The latter two clusters show a Th2-oriented profile, as indicated by *ZBTB16* and *GATA3* expression (**Figure 3E**).

After expansion, three additional clusters were identified, characterized by predominant expression of cell cycle–associated genes and corresponding to cells in the S, G1, and G2/M phases (**Figure 3A-C-D**). They contain both CD4^+^ and CD4^-^ populations, are not activated (CD69 neg) neither mature with absence of CD161 (KLRB1) expression (**Figure 3B-D**).

Cluster distributions changed markedly between day 0 and day 14, indicating distinct expansion dynamics across iNKT subsets. While the cycling clusters were only observed on day 14, the ‘CD4^+^ IFIT2’ cluster was detected only on day 0, and the proportions of the ‘CD4^+^ LEF1’, ‘CD4^−^ SLC4A10’, and ‘CD8^+^’ clusters declined over time. By contrast, the ‘CD4^−^ GNLY’ and ‘CD4^−^ XCL1’ populations increased after expansion with both cytokines (**Figure 3F**).

We next performed pseudotime and CytoTRACE analyses to define the differentiation trajectories of the expanded iNKT subsets (**Figure 4**). The ‘CD4^−^ CD247’ cluster emerged as the least differentiated population and the one most closely related to the G2/M cluster, consistent with shared transcriptomic features including high *CD247* expression (**Figures 3C-E and 4**). Gene Ontology analysis indicated enrichment for protein refolding and cell structure-related processes in this cluster (**Figure 4B**). Trajectory analysis further suggests differentiation toward either CD4^+^ or CD4^−^ subsets (**Figure 4A**). Among CD4^+^ cells, the rare ‘CD4^+^ LEF1’ population was associated with post-thymic differentiation and proliferation pathways, whereas the major CD4^+^ subset showed enrichment for proliferation, cytokine production, and antiviral response programs (**Figures 4C-D**). Among CD4^−^ cells, the predominant ‘CD4^−^ SLC4A10’ subset was associated with T-cell activation, Th polarization, and differentiation pathways, whereas the ‘CD4^−^ GNLY’, ‘CD4^−^ XCL1’, and CD8^+^ subsets appeared more differentiated and were enriched for cytotoxic functions; the first two also showed signatures related to chemotaxis and migration (**Figures 4F-H**).

**Figure 4:**
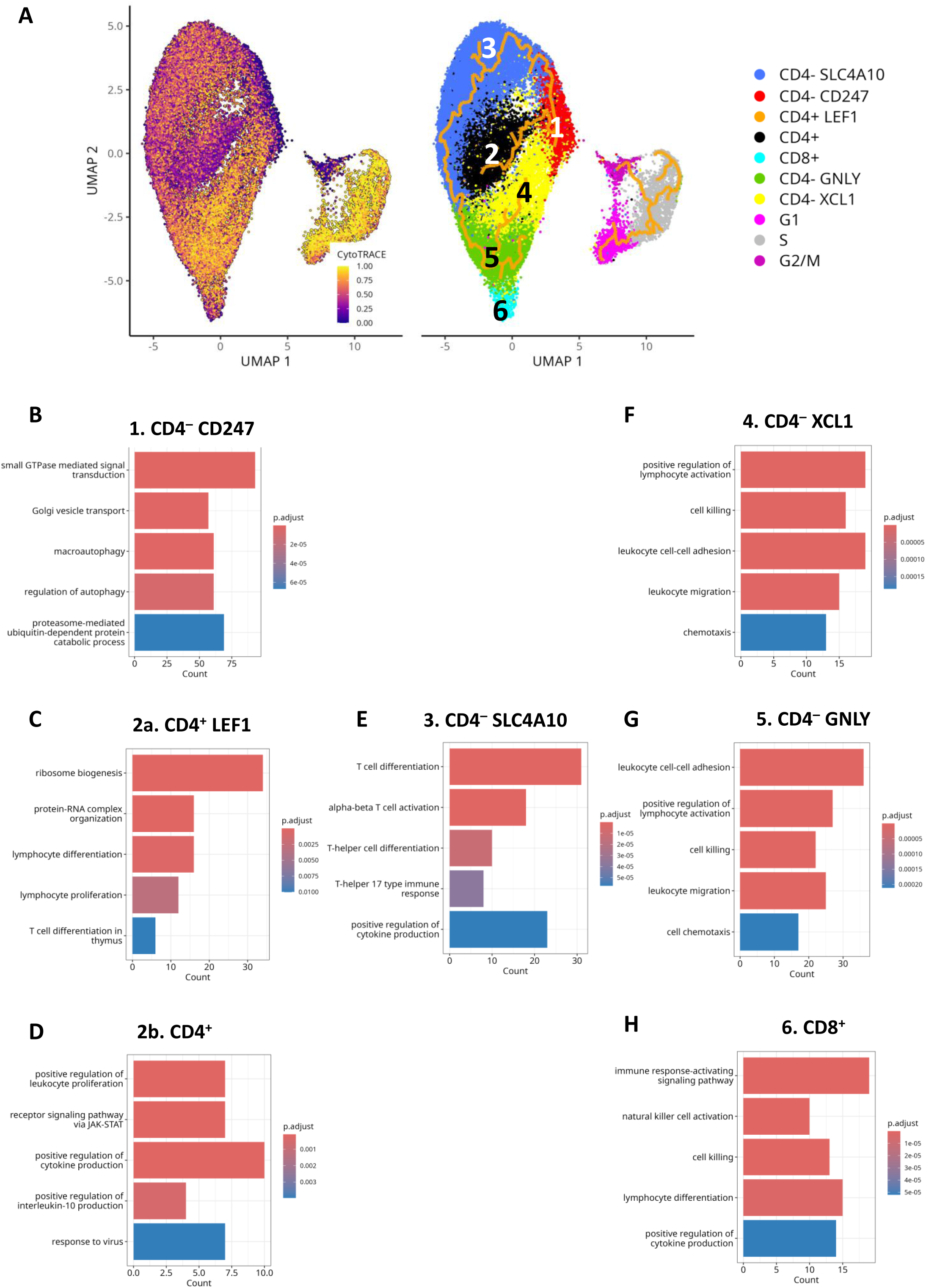
Developmental trajectories and characteristics of iNKT cell clusters. (A) CytoTRACE prediction and trajectory analysis of the single-cell RNA-seq iNKT dataset. (B– H) Bar plots showing enriched Gene Ontology biological process terms among genes significantly upregulated in each cluster, excluding cell cycle-related clusters.

### Comparison between IL-15 and IL-2 expanded iNKT cells

We then compared genes upregulated in iNKT cells expanded with IL-15 versus IL-2, and vice versa (**Figure 5A**). Functional annotation of genes enriched after IL-2 expansion indicated a profile associated with immune responses, particularly antiviral pathways. These cells also showed a proliferative phenotype and a transcriptomic metabolic signature consistent with glycolytic and cytotoxic functions (**Figure 5B**). By contrast, IL-15 expansion was associated with upregulation of genes involved in T-cell differentiation, strong transcriptional activity, and TGF-β responsiveness. These cells also showed gene programs related to cytokine regulation and modulation of immune and inflammatory responses (**Figure 5C**).

**Figure 5:**
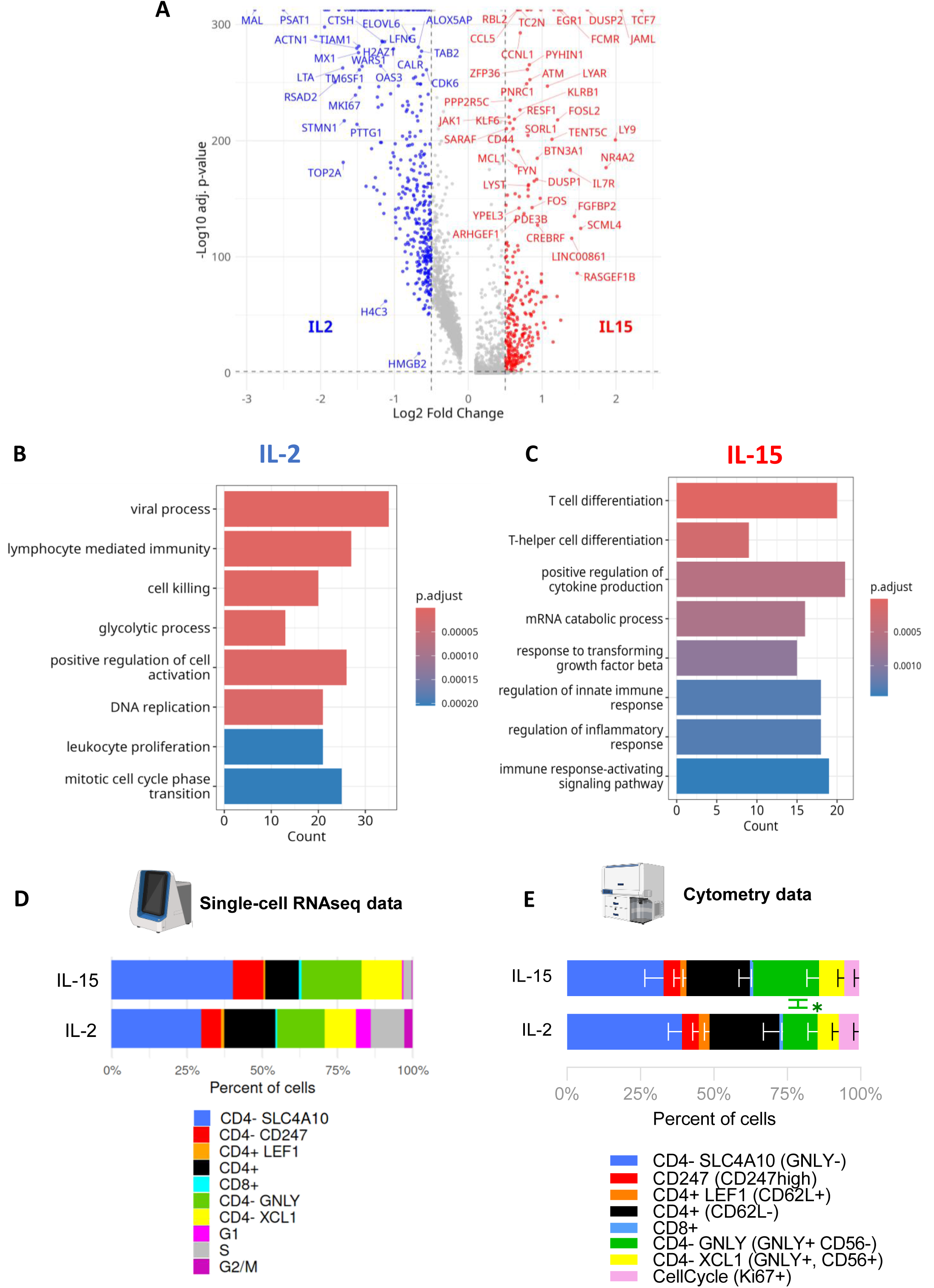
Transcriptomic and cytometric profiling highlights enhanced differentiation and cytotoxicity in IL-15-expanded iNKT cells compared with IL-2-expanded cells. (A) Volcano plot of differentially expressed genes between IL-15 and IL-2 conditions. Significantly different genes are shown in red based on a t-test accounting for inter-sample variability. Enriched Gene Ontology biological process terms for genes significantly upregulated across samples are shown for (B) IL-2 and (C) IL-15. Bar plots show the distribution of iNKT-cell clusters after expansion with IL-15 or IL-2, as assessed by (D) single-cell RNA-seq and (E) flow cytometry. Statistical comparisons were performed using a mixed-effects model with Šídák’s multiple-comparisons test.

At the transcriptomic level, IL-2 preferentially enriched cells in S phase (*p*=0.021), with a similar trend for G1 (*p*=0.061) (**Suppl. Figure. 3**) and tended to increase the proportions of ‘CD4^+^’ and ‘CD4^+^ LEF1’ clusters. By contrast, IL-15 significantly increased the proportions of the ‘CD4^−^ GNLY’ (*p*=0.011) and ‘CD4^−^ XCL1’ (*p*=0.016) populations (**Suppl. Figure 3**). Flow cytometry using the gating strategy shown in **Suppl. Figure 4** confirmed the identification of clusters defined by transcriptomic analyses (**Figure 5E**). Phenotypic analysis further showed that IL-15 expansion generated a higher proportion of ‘CD4^−^ GNLY’ cells than IL-2 expansion (*p*=0.0489) (**Figure 5E**).

### Control of GvHD by expanded iNKT cells

We next assessed the functional properties of iNKT cells expanded with the optimized IL-15 protocol. After expansion, iNKT cells can produce IFN-γ upon stimulation (**Figure 6A**), express high levels of perforin and granzyme B (**Figure 6B**). They also express NK activating NKG2D and inhibitory NKG2A molecules at similar levels (**Figure 6C**). These features were similar in total iNKT cells expanded with IL-2 or IL-15 (**Suppl. Figure 5**).

**Figure 6:**
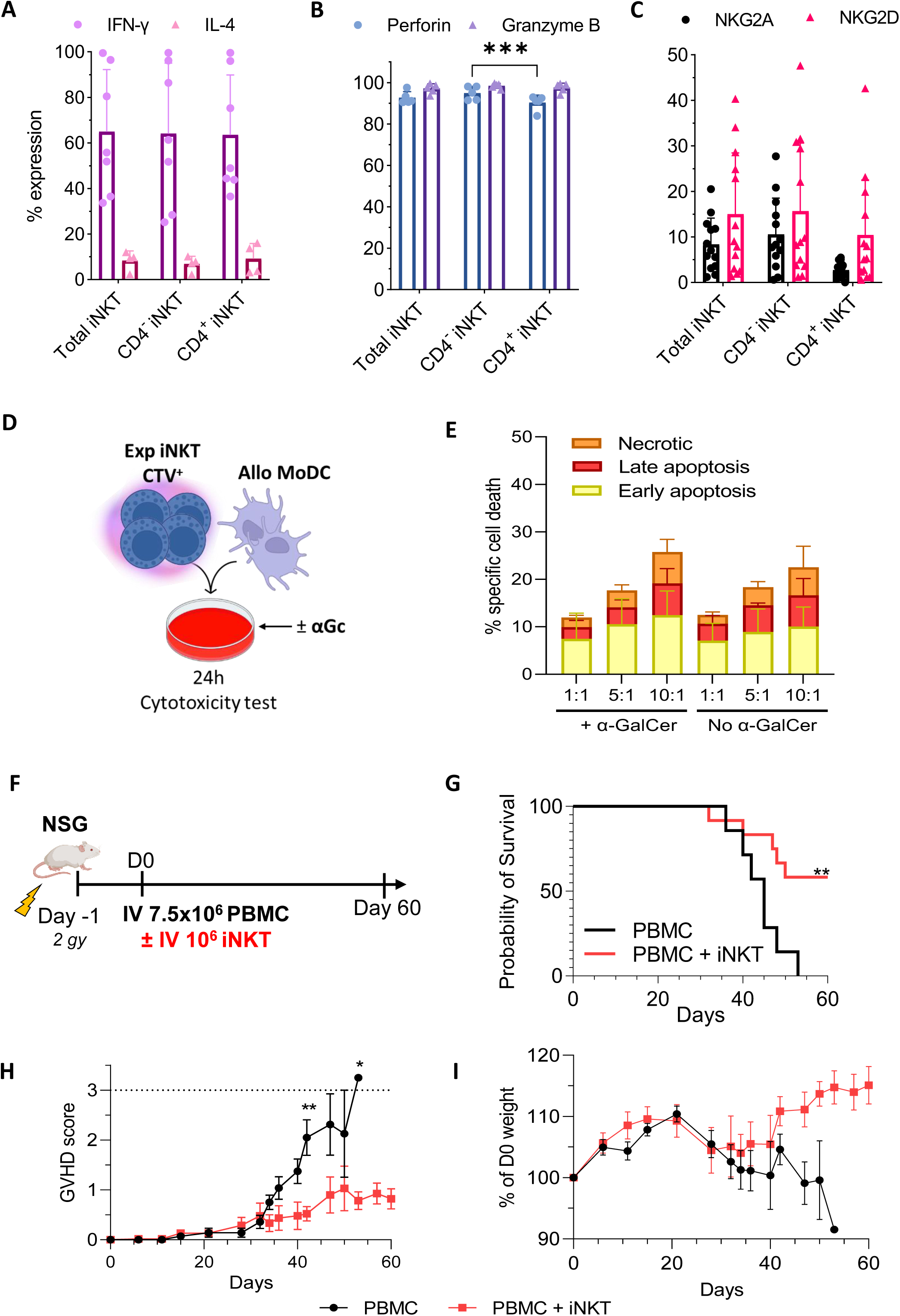
Expanded iNKT cells retain functional activity and suppress GvHD in vitro and in vivo. (A) Expression of IFN-γ (n=7) and IL-4 (n=4) after 4 h of PMA/ionomycin stimulation in total, CD4−, and CD4+ expanded iNKT cells. Data are shown as mean + SD. (B) Expression of perforin and granzyme B in total, CD4−, and CD4+ iNKT cells after expansion (n=6). Data are shown as mean + SD; comparisons were performed using two-way ANOVA. (C) Expression of NKG2A and NKG2D in total, CD4−, and CD4+ iNKT cells after expansion (n=5). Data are shown as mean + SD; comparisons were performed using two-way ANOVA. (D) Schematic of the 24-hour cytotoxicity assay using expanded iNKT cells labelled with CellTrace Violet (CTV) and MoDCs cultured with or without α-GalCer. (E) Representative flow cytometry plots showing Annexin V and 7-AAD staining in dendritic cells after 24 h of co-culture with iNKT cells. Early apoptosis (Annexin V+, yellow), late apoptosis (Annexin V+7-AAD+, red), and necrosis (7-AAD+, orange) are indicated. (F) Induction of early apoptosis (yellow), late apoptosis (red), and necrosis (orange) in dendritic cells after 24 h of co-culture with expanded iNKT cells at different effector-to-target (E:T) ratios, with or without α-GalCer (n=3). Data are shown as mean + SEM. (G) Schematic of the in vivo anti-GvHD study in NSG mice. Mice were irradiated at 2 Gy 24 h before injection of 7.5×10^6^ human PBMCs, with or without 10^6^ expanded human iNKT cells. (H) Kaplan-Meier survival curves of NSG mice after GvHD induction without (n=7) or with (n=12) supplementation of 10^6^ total iNKT cells expanded for 14 days. (I) Clinical GvHD scores and (J) body weight relative to D0 over 60 days in the PBMC group (black, n=7) and the PBMC+iNKT group (red, n=12). Data are shown as mean ± SEM; comparisons were performed using multiple t-tests.

Because, we previously showed that IL-2-expanded CD4^−^ iNKT cells can control graft-versus-host disease (GvHD) by lysing dendritic cells (DCs)^13^, we performed 24-hour cytotoxicity assays using predominantly CD4^−^ IL-15-expanded iNKT cells and allogeneic monocyte-derived dendritic cells (Allo-MoDCs) at different effector-to-target (E:T) ratios, with or without α-GalCer (**Figure 5D**). After 24 hours of co-culture with iNKT cells, MoDCs were stained with Annexin V and 7-AAD to distinguish early apoptosis (Annexin V^+^7-AAD^−^), late apoptosis (Annexin V^+^7-AAD^+^), and necrosis (Annexin V^−^7-AAD^+^) within the CTV-negative population. iNKT cells generated with the optimized expansion protocol induced MoDC death in a dose-dependent manner (**Figure 5E**). At an effector-to-target ratio of 10:1, the mean (SD) percentage of apoptotic or necrotic MoDCs was 25.76% (SD 6.23%) in the presence of α-GalCer and 22.53% (SD 6.99%) without α-GalCer, with no significant difference between conditions (*p*=0.9603).

We next evaluated the ability of IL-15-expanded iNKT cells to control GvHD in a xeno-GvHD model using immunodeficient NSG mice engrafted with human PBMCs. Mice were irradiated one day before transplantation and received PBMCs either alone or supplemented with autologous total iNKT cells. Survival, clinical GvHD score, and body weight were monitored for 80 days (**Figure 5F**). Administration of 10^6^ IL-15-expanded iNKT cells significantly prolonged mouse survival compared with the GvHD control group (median survival not reached vs 45 days, respectively; *p*=0.0078) (**Figure 5G**). Relative to the PBMC-only group, mice receiving iNKT cells (PBMC+iNKT) showed lower clinical GvHD scores from day 42 onward (*p*=0.0032) and a trend toward reduced weight loss from day 47 (*p*=0.1298) (**Figure 5H-I**). GvHD control was also maintained at a lower iNKT dose, of 2.5×10^5^ iNKT cells for 7.5×10^6^ PBMCs (**Suppl. Figure S6**).

### Preservation of GvL effect by expanded iNKT cells

To assess whether GvHD control could be achieved without impairing, and potentially while enhancing, the graft-versus-leukaemia (GvL) effect of hematopoietic stem cell transplantation (HSCT), we evaluated the cytotoxic activity of IL-15-expanded iNKT cells against human leukaemic cells. We first performed in vitro cytotoxicity assays using the CD1d-expressing leukaemia cell lines THP-1 (myeloid) and Jurkat (lymphoid). Expanded iNKT cells were co-cultured for 16 hours with THP-1 or Jurkat cells at different effector-to-target (E:T) ratios, with or without α-GalCer (**Figure 7A**). iNKT cells lysed both leukaemic cell lines. For THP-1 cells, specific lysis at a 1:1 ratio was similar in the presence and absence of α-GalCer (mean [SD], 34.75% [15.87] vs 24.03% [17.99]; *p*=0.4594). In contrast, Jurkat cell lysis was strictly α-GalCer-dependent (mean [SD] at 1:1, 33.57% [18.40] with α-GalCer vs 7.21% [8.62] without; *p*=0.0014) (**Figure 7B**). Total IL-2-expanded iNKT cells showed similar anti-leukaemic activity in vitro (**Suppl. Figure 7**).

**Figure 7:**
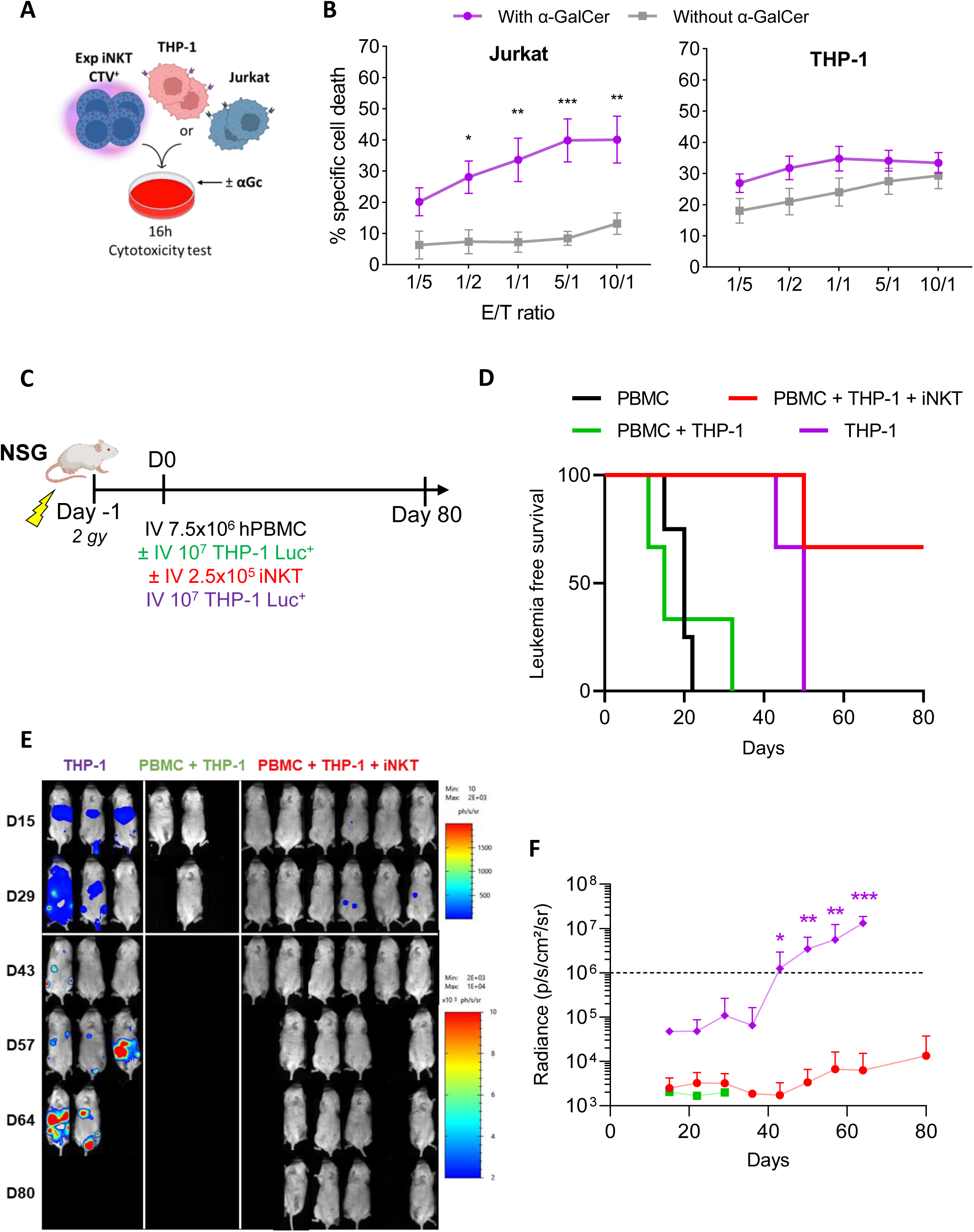
Expanded iNKT cells show robust anti-leukemic activity in vitro and in vivo in a combined GvL/GvHD model. (A) Schematic of the 16-hour cytotoxicity assay using expanded iNKT cells labelled with CellTrace Violet (CTV) and the leukaemia cell lines THP-1 or Jurkat, with or without α-GalCer. (B) Specific tumour cell death in Jurkat cells (left panel, n=6) and THP-1 cells (right panel, n=8), with or without α-GalCer. Data are shown as mean ± SEM; comparisons were performed using two-way ANOVA with Šídák’s test. (C) Schematic of the in vivo study of GvL and anti-GvHD activity in NSG mice. Mice were irradiated at 2 Gy 24 h before injection of 7.5×10^6^ human PBMCs, with or without 10^7^ THP-1 leukaemia cells, and with or without 2.5×10^5^ expanded human iNKT cells. Clinical GvHD score, body weight, and leukemia progression by bioluminescence were monitored for 80 days. (D) Kaplan-Meier curve of leukemia-free survival, defined as bioluminescence lower than 10^6^ ph/s/cm²/sr, in NSG mice after induction of the GvHD effect (PBMC, n=4), leukemic cells alone (THP-1, n=3), GvL/GvHD (PBMC+THP-1, n=3), or GvL/GvHD supplemented with expanded iNKT cells (PBMC+THP-1+iNKT, n=6). (E) Bioluminescence imaging of tumor progression in groups receiving THP-1 cells. (F) Radiance over time according to the experimental group.

We then assessed whether IL-15-expanded iNKT cells could simultaneously control GvHD and preserve GvL activity in a xeno-GvHD/GvL mouse model. Immunodeficient NSG mice were irradiated one day before cell transfer and then injected with human PBMCs, either alone or together with autologous total iNKT cells and/or Luc^+^ THP-1 cells (**Figure 6C**). All mice that received PBMCs without iNKT cells, with or without THP-1 cells, died of GvHD by day 35 (**Figure 6D-E**). Mice injected with THP-1 cells alone developed significant leukaemia progression after day 40 (**Figure 6F**). In the PBMC+THP-1 group, mice died from GvHD before developing detectable tumor burden. By contrast, mice receiving PBMC+THP- 1+iNKT cells showed prolonged survival, with more than 60% remaining alive at the end of the 80-day study period, together with low bioluminescence signals and low GvHD scores, indicating concurrent leukemic control and GvHD protection (**Figure 6D-F and Suppl Figure 8**).

## Discussion

iNKT-cell frequencies vary widely among individuals, ranging from 0.001% to 1% of T lymphocytes. Their use in cellular immunotherapy therefore requires clinical-grade protocols are needed to expand these cells, particularly the CD4^−^ subset. Among the cytokines tested for T-cell expansion (IL-2, IL-4, IL- 7, and IL-15) ^26^, IL-2 and IL-15 were the most effective for iNKT-cell proliferation, in line with previous comparisons of IL-2, IL-7, IL-12, and IL-15 ^27,28,17,20^. We showed that IL-15 alone achieved the strongest expansion of fully differentiated and cytotoxic CD4^−^ iNKT subpopulations compared with the other cytokines tested. These results support previous observations by Metelitsa and colleagues showing selective expansion of CD4^−^ iNKT cells induced by IL-15 ^29^. This effect has been linked to higher surface expression of CD122, the shared beta-chain of the IL-2 and IL-15 receptors, whereas CD4^+^ iNKT cells preferentially express CD127, the IL-7 receptor ^29^. Consistent with this, IL-7 increased the proportion of CD4^+^ iNKT cells in our cultures but did not induce substantial iNKT-cell proliferation. Finally, our phenotypic and transcriptomic analyses confirmed that IL-2 preferentially supports CD4^+^ iNKT cells, consistent with established iNKT-cell expansion protocols ^16–20^. To our knowledge, our method represents the first approach enabling rapid, high-level expansion of human CD4^−^ iNKT cells, achieving 100- to 10,000-fold expansion within 14 days. The protocol is compatible with GMP culture conditions and is expected to generate more than 2×10^8^ CD4^−^ iNKT cells from a lymphapheresis containing at least 10^9^ PBMCs. Although this protocol generates fewer cells than prolonged IL-2-based expansion methods ^16,30^, its shorter duration may limit cellular exhaustion and promote greater proliferative capacity *in vivo*.

To better define our cellular therapy product, we performed single-cell RNA sequencing of iNKT cells before and after expansion. Analysis of baseline cells also allowed us to contribute to the characterization of circulating human iNKT-cell subsets, a field that remains poorly documented by single-cell transcriptomic studies. The first single-cell RNA-seq analysis of human peripheral blood iNKT cells compared sorted unstimulated cells with PMA/ionomycin-stimulated cells and identified four major subsets but this study was limited by the relatively small number of unstimulated iNKT cells analyzed and few immunological and functional characterization of the identified subsets ^31^. Using the same single-cell platform than we used, analysis of juvenile human thymic iNKT cells identified markers associated with progressive maturation, including SOX4, inversely related to maturity, and KLRB1, which increases with cellular maturation^32^. More recently, the same group extended this analysis to iNKT cells from human thymus, cord blood, bone marrow, and peripheral blood, revealing tissue-specific subsets in thymus and cord blood and greater subset diversity in peripheral blood and bone marrow^33^. Several CD4^−^ peripheral blood populations described in that study were also detected in our dataset, including CD4^+^ IFIT2, CD4^+^ LEF1, CD4^−^ GNLY, and CD4^−^ CD247 subsets. These comparisons support the robustness of the main human iNKT-cell populations identified in our study while highlighting the need for harmonized analytical frameworks to refine the classification of human iNKT subsets. These convergent studies also highlight major differences between mouse and human iNKT-cell subsets, particularly regarding Th1, Th2, and Th17 differentiation. Whereas mouse iNKT subsets display distinct transcriptomic programs defined by Tbx21, Rorc, and Zbtb16 expression ^34^, we and others ^33^ could not identify such clearly segregated profiles in human iNKT cells.

In addition, our study provides new insight into the characteristics of iNKT cells expanded with IL-2 or IL-15. Although overall iNKT-cell diversity was preserved after expansion, subset composition changed substantially, with loss of the ‘CD4+ IFIT2’ cluster, marked reduction of the ‘CD4+ LEF1’ cluster, and strong expansion of the ‘CD4^−^ GNLY’ and ‘CD4^−^ XCL1’ subsets. The reduction of the ‘CD4^+^ LEF1’ cluster may reflect its immature, stem-like phenotype, characterized by *SELL* (CD62L) and *LEF1* expression ^35,36^. These cells may respond to cytokine stimulation but rapidly differentiate after TCR engagement by α- GalCer during expansion, similar to the transition of stem cell memory T cells toward central memory T cells after antigenic stimulation in the presence of IL-7 and IL-15 ^37^. This stem-like iNKT state has been shown to be attractive for CAR-iNKT strategies because of its potential for long-term persistence ^36,38^, but this expansion appears to require IL-21 in combination with IL-2 ^38^. The ‘CD4^+^ IFIT2’ iNKT cluster likely represents cells recently activated by type I interferons, possibly in response to viral infection ^39^. Without sustained interferon signaling during *in vitro* expansion, this transcriptional program may not be maintained. A similar cluster has also been reported by Mavers’ group in both peripheral blood and cord blood ^33^.

Cytokine stimulation induced cell cycling and promoted expansion of CD4^+^ and most CD4^−^ subsets, with the exception of the predominant CD4^−^ SLC4A10 population. Notably, this gene was recently shown to be upregulated in adult iNKT cells compared with cord blood iNKT cells in a study by Trujillo-Ocampo et al. ^15^. Trajectory analyses suggested that cycling cells transition through an intermediate CD4^−^ CD247^+^ stage, enriched in CD4^−^ iNKT cells, before diverging toward either CD4^+^ or CD4^−^ lineages. Compared with CD4^−^ subsets, CD4^+^ iNKT cells appeared less mature and less differentiated, with the CD4^+^ LEF1 cluster representing the least differentiated population. Gene Ontology analyses support this interpretation, showing for the latter features of post-thymic differentiation, high proliferative and differentiation potential, and limited migratory or cytotoxic activity ^32,33^. The broader CD4^+^ compartment also retained strong proliferative capacity, cytokine-production programs, and antiviral-response pathways. In contrast, the CD4^−^ compartment followed a more advanced differentiation trajectory toward cytotoxic effector states. This lineage appeared to arise from the large CD4^−^ SLC4A10^+^ pool of activated, differentiating iNKT cells towards three terminal CD4^−^ clusters: CD4^−^ XCL1, CD4^−^ GNLY, and CD8^+^, characterized by progressively greater differentiation, migratory capacity, and cytotoxic potentials.

Our comparative analysis revealed distinct functional programs induced by IL-2 and IL-15 during iNKT-cell expansion. IL-2 preferentially promoted proliferating iNKT cells with strong antiviral gene signatures, consistent with the enrichment of CD4^+^ subsets and cycling cells. This antiviral profile may explain the reported efficacy of IL-2-expanded allogeneic iNKT cells in ARDS ^40^. In contrast, IL-15 favored iNKT-cell differentiation and immunoregulatory programs, together with enrichment of terminally differentiated CD4^−^ XCL1 and CD4^−^ GNLY subsets. These populations display strong migratory and cytotoxic features that may contribute to GvHD control while preserving a potent GvL effect in the allo-HSCT setting. Despite marked phenotypic and transcriptomic differences, bulk iNKT-cell products expanded with IL-2 or IL-15 showed comparable cytotoxicity activity *in vitro*. This shared potency likely reflects the persistence, within IL-2-expanded cultures, of all major iNKT subsets, including highly cytotoxic populations.

Finally, our study confirms that expanded human iNKT cells can control GvHD in a conventional xeno-GvHD model, as previously reported by our group and others ^13,15,41^. Importantly, we show for the first time that IL-15-expanded total human iNKT cells, enriched in CD4^−^ iNKT cells, can both limit GvHD and preserve GvL activity in a preclinical GvHD/GvL mouse model. In mice, iNKT2 (Zbtb16^+^) and iNKT17 (Rorc^+^) subsets have been shown to protect against GvHD, whereas iNKT1 (Tbx21^+^) cells exert the strongest GvL effect ^15,34^. Our data indicates that IL-15-expanded human iNKT cells, which include all Th-related subsets, can both control GvHD and preserve GvL activity.

Several mechanisms may account for iNKT-mediated GVHD control. In mice, iNKT2 and iNKT10 cells promote regulatory T-cell induction ^15,42^, whereas our previous work showed that human IL-2-expanded CD4^−^ iNKT cells modulate antigen-presenting cells (APCs), thereby limiting allogeneic T-cell activation^13^. Our in vitro assays suggest that IL-15-expanded human iNKT cells, enriched in differentiated and highly cytotoxic Th1-like subsets, may further enhance leukemic cell targeting in addition to regulating APCs. This feature may represent an advantage over cord blood-derived expanded human iNKT cells, which are enriched in CD4^+^ IL-10-producing cells with suppressive but limited cytotoxic properties ^15^. Although other cellular therapies, including mesenchymal stromal cells ^43^ and regulatory T cells ^44–46^, have been proposed for GvHD prevention, they do not provide direct anti-leukemic or antiviral activity. Interestingly, iNKT cells may also be effective at low iNKT-to-T-cell ratios. Clinically, a peripheral blood ratio greater than 1 iNKT cell per 1,000 T cells at day 15 after HSCT was associated with an 80% probability of remaining GvHD-free, compared with 19.5% below this threshold ^12^. Based on our in vivo data, in which an iNKT:T-cell ratio of 1:15 reduced xeno-GvHD, and on the typical HSCT graft content of 1–5×10^7^ T cells/kg, we estimate that 1–3×10^6^ iNKT cells/kg may be sufficient for a phase I dose-escalation trial. Allogeneic iNKT cells have already been evaluated in phase I trials for viral-induced ARDS and solid tumors, with maximum tested doses of 10^9^ total iNKT cells and 1.4×10^7^ iNKT cells/kg, respectively ^47,40^. Published data indicate a favorable safety profile, with no reported cytokine release syndrome or neurotoxicity ^47,40,48^.

In summary, we report a rapid, scalable, GMP-compatible approach to generate clinically relevant numbers of CD4^−^ iNKT cells directly from total PBMCs. By integrating single and functional analyses in preclinical models, we show that IL-15-driven expansion preserves iNKT-cell diversity while driving cytotoxic CD4^−^ effector programs. By coupling GvHD control without impairing anti-leukemic activity, these data support IL-15-expanded iNKT cells as a promising compelling immunotherapy candidate for allo-HSCT.

## Materials and Methods

### In vitro cultures

Human iNKT cells were expanded from PBMCs obtained from healthy volunteers through the Etablissement Français du Sang (EFS) (n°L20DIV0856). PBMCs were cultured for 14 days with 100 ng/mL alpha-galactosylceramide (α-GalCer; KRN7000, Funakoshi). The cytokines rhIL-2 IS (845 IU/mL, Miltenyi Biotech), rhIL-15 (5-50 ng/mL, Miltenyi Biotech), rhIL-4 (10 ng/mL, Miltenyi Biotech), and rhIL- 7 (10 ng/mL, Miltenyi Biotech) were added on day 0. When indicated, 50% of the medium was replaced on day 7 and cytokines were re-added. For the boost condition, PBMCs from the same donor were added on day 14 at the same number as on day 0, together with cytokines and α-GalCer at the initial concentrations. Cell density varied according to the culture format: 5×10^5^ cells/mL for 2 mL cultures in 24-well plates and 100 mL culture bags (Miltenyi Biotech), and 2×10^6^ cells/cm^2^ (2×10^5^ cells/mL) in 100 mL 6M G-Rex bioreactors (Wilson-Wolf). RPMI 1640 (Gibco) was supplemented with 2 mM GlutaMAX, 25 mM HEPES, 10% FCS (Dutscher/Pan Biotech/Hyclone), and 1% penicillin-streptomycin (Sigma-Aldrich) was used for iNKT, MoDC, and leukaemia cell (THP-1 and Jurkat) cultures. RPMI 1640 Advanced medium (Gibco) was supplemented with sodium pyruvate, non-essential amino acids, 5% FCS (Dutscher/Pan Biotech/Hyclone), and 4 mM glutamine was used as the optimal iNKT culture medium. Cultures were maintained at 37°C in a normoxic incubator with 5% CO_2_. Lactate and glucose concentrations in culture supernatants were measured using a StatStrip Xpress 2 analyzer (NOVA Biomedical) with dedicated lactate and glucose test strips.

### MoDC generation

CD14^+^ cells were positively isolated using CD14 microbeads (Miltenyi Biotech) and cultured in complete RPMI supplemented with IL-4 (20 ng/mL, Miltenyi Biotech) and GM-CSF (100 ng/mL, Miltenyi Biotech) for 5 days. Immature MoDCs were then cultured for an additional 2 days in complete RPMI containing TNFα (50 ng/mL, Miltenyi Biotech) and PGE2 (1 µg/mL, Focus Biomolecules) to generate mature MoDCs.

### Flow cytometry

Surface staining of cultured cells was performed for 20 min at 4°C in PBS containing 0.5% BSA. Antibodies are listed in **Suppl. Table 1**. Cells were analyzed on a Celesta Sorp (BD Biosciences) or a Gallios cytometer (Beckman Coulter) using Kaluza software (Beckman Coulter). iNKT cells were identified as CD3^+^ cells co-expressing the invariant iNKT T-cell receptor (iTCR), and were quantified as a percentage of total T lymphocytes and in absolute numbers. For intracellular cytokine staining (IFN-γ and IL-4), cells were stimulated for 4 h in complete RPMI with cell stimulation cocktail and protein transport inhibitors (eBioscience) at 37°C and 5% CO_2_. Matched control samples were incubated with protein transport inhibitors alone. Cells were first stained with a fixable viability dye, then processed for intracellular staining using the Foxp3 Fix/Perm kit (eBioscience) according to the manufacturer’s instructions.

### Single-cell RNAseq preparation, capture and sequencing

Before expansion (D0) or after 14 days of expansion (D14), iNKT cells were isolated using anti-iNKT microbeads (Miltenyi Biotech). Non-expanded D0 samples were sorted twice. Each D14 sample was generated under two conditions, IL-2 and IL-15. Cells were co-stained with Sample-Tag for multiplexing and with an Ab-Seq panel (CD3, CD4, CD8, CD161, CD56, and iNKT TCR) to assess surface protein expression. Staining, cell capture, and library preparation for whole-transcriptome analysis (WTA), Ab-Seq, and Sample-Tag were performed according to the BD Rhapsody protocol. Libraries were sequenced on a NovaSeq 6000 (Illumina) at the Imagine Institute facility (Necker Hospital, Paris) with a depth of 50,000 reads per cell.

### Data processing and integration workflow

Demultiplexing was performed using the BD Genomics pipeline, and the resulting DBSEQ files were analyzed in R with Seurat (v5). Cells were filtered using standard quality control criteria applied uniformly across samples: nFeature_RNA > 200, nFeature_RNA < 5000, and mitochondrial content < 25%. Ab-Seq features were retained in the RNA slot for dimensionality reduction and clustering

because this improved robustness and were also stored in the ADT slot after CLR normalization for complementary analyses. Log-transformed CD3 and NKT TCR Ab-Seq histograms showed clear bimodal distributions with a trough around 3, allowing iNKT-cell selection using thresholds of CD3 > 3 and NKT TCR > 3. Only cells meeting these criteria were included in the integrated dataset. RNA data were log-normalized and integrated across samples using Seurat’s CCA-based workflow (SelectIntegrationFeatures, FindIntegrationAnchors, and IntegrateData). Dimensionality reduction was performed using PCA and UMAP on the integrated RNA assay, and both RNA and protein data were used for downstream clustering and annotation.

### Cytotoxicity test

iNKT cells were isolated using anti-iNKT microbeads (Miltenyi Biotech), stained with CellTrace Violet (Life Technologies), and co-cultured with either mature MoDCs for 24 h or the leukaemia cell lines THP- 1 and Jurkat for 16 h in target medium at different effector-to-target ratios. Apoptosis and necrosis were assessed by 7-AAD and Annexin V-FITC staining (Sony Biotechnology) in 1X Annexin V buffer (Sony Biotechnology). Specific cytotoxicity was calculated as ((% sample (7-AAD^+^, CTV^−^) − % spontaneous (7- AAD^+^, CTV^−^)) / (100 − % spontaneous (7-AAD^+^, CTV^−^))) x 100. All assays were performed in duplicate or triplicate.

### Animal experiments

Female NSG mice aged 8–12 weeks from the local animal facility (ACBS, University of Lorraine) received 2 Gy total-body irradiation (X-RAD320, CRAN Lab facility) on day −1. On day 0, mice were transplanted with 7.5×10^6^ human PBMCs either alone (PBMC group) or together with 10^6^ human iNKT cells expanded as described above from the same donor. Mice were monitored twice weekly for body weight, survival, and clinical GvHD score, as previously described^13^. In mice receiving luciferase-positive THP-1 cells (BPS Bioscience), tumour burden was assessed weekly by bioluminescence imaging using a PHOTON Imager Optima (Biospace Lab) after intraperitoneal injection of D-luciferin (150 mg/kg; Promega). All animal procedures were approved by the French Ministry of Higher Education and Research and by the local ethics committee in Nancy, the ‘Comité d’Ethique Lorrain en Matière d’Expérimentation Animale’ (CELMEA), under protocol APAFIS#40977.

### Statistics

Statistical test use and number of samples are specified in legend of appropriate figure or shown on the figures. Figures were created and analyzed with GraphPad Prism 9.0.0. For all statistical tests, p- values are identified as follows: ∗p < 0.05, ∗∗p < 0.01, ∗∗∗p < 0.001, and ∗∗∗∗p < 0.0001.

## Data availability statement

Data supporting the findings of this study are available from the corresponding author upon reasonable request. Single-cell data are available at GEO under accession number GSE330507.

## Acknowledgments

This work was supported by the French National Cancer Institute (INCa), and SATT SAYENS. This project has benefited from a CPER Grand Est I2GE 2021-2027. We thank Huguette Louis from the Cytometry Core Facility of Shared Platform Service SMP IBSLor (Université de Lorraine). We thank Joël Daouk from the irradiation platform of the CRAN laboratory (Université de Lorraine), as well as the central animal facility team of the University of Lorraine (ACBS). Sequencing analyses were carried out by the sequencing platform at the Imagine Institute in Paris.

## Author contributions

J.B. and MT.R. conceptualized the project. J.B., C.C., L.M., G.F., S.H., S.P., D.M. and MT.R. collected and analyzed the data. MT.R. supervised the project. J.B., C.C., T.C. and MT.R. wrote the manuscript. All authors read and approved the final manuscript.

