## Supplementary data for "Scalable expansion of human iNKT cells: single-cell profiling and in vivo control of GvHD with preserved GvL activity"

**Supplementary Table 1: List of antibodies and reagents used for flow cytometry**

| Target | Fluorochrome | Clone | Manufacturer |
|---|---|---|---|
| **CD3** | FITC | OKT3 | Sony |
| **CD3** | PE | OKT3 | Sony |
| **CD3** | AF700 | UCHT1 | BD |
| **CD4** | BV421 | RPA-T4 | Sony |
| **CD4** | PE-Cy7 | OKT4 | Sony |
| **CD4** | PE-Cy5 | S3.5 | Invitrogen |
| **CD8** | PerCp-Cy5.5 | SK1 | BD |
| **TCR Vα24-Jα18 (iNKT)** | BV750 | 6B11 | BD |
| **TCR Vα24-Jα18 (iNKT)** | PE | REA1054 \| 6B11 | Miltenyi |
| **TCR Vα24-Jα18 (iNKT)** | APC | 6B11 | Miltenyi |
| **NKG2D** | BV510 | 1D11 | Sony |
| **NKG2A** | PE-CF594 | S19004C | Sony |
| **IFNγ** | BV421 | B27 | Sony |
| **IL-4** | PE | MP4-25D2 | Sony |
| **Perforin** | PE | dG9 | Sony |
| **Granzyme B** | PE-Dazzle 594 | QA16A02 | Sony |
| **CD161** | BV421 | HP-3G10 | Sony |
| **PLZF** | PE | Mags.21F7 | Invitrogen |
| **TBET** | BV786 | O4-46 | BD |
| **RORC** | PE-CF594 | Q21-559 | BD |
| **CD56** | APC-Cy7 | 5.1H11 | Sony |
| **CD56** | PE | HCD56 | Sony |
| **CD247** | AF647 | 6B10.2 | BD |
| **CD62L** | BV711 | DREG-56 | Sony |
| **GNLY** | AF488 | RB1 | BD |
| **Ki67** | BV605 | KI-67 | Sony |
| **CD69** | PE-Cy7 | PE-Cy7 | Sony |
| **Viability** | DY506 | / | BD |

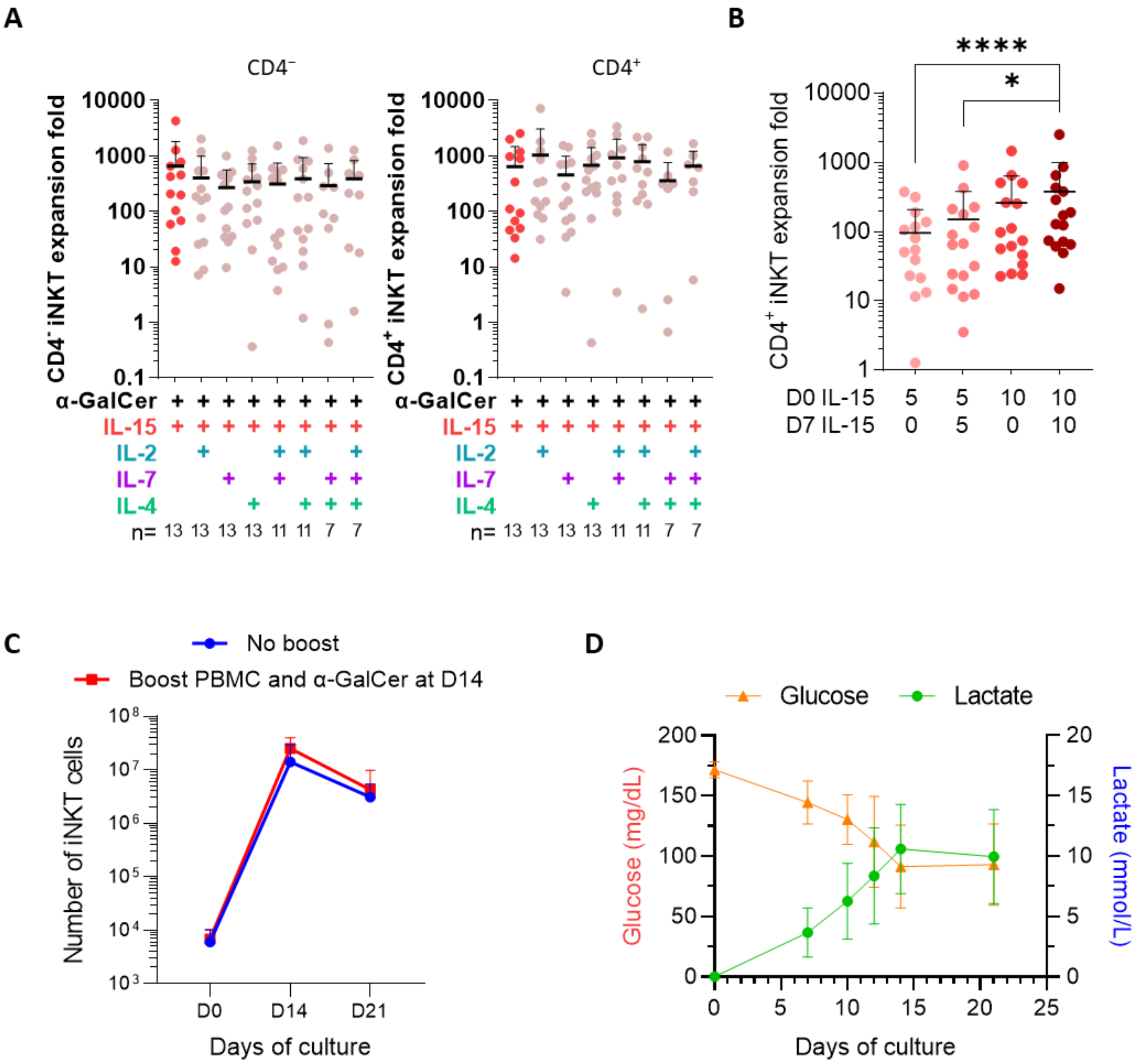


**Supplementary Figure 1: Alternative cytokine combinations and culture conditions for iNKT cell expansion.**

(A) Expansion of CD4− and CD4+ iNKT cells over 14 days under different cytokine regimens: IL-15 alone or combined with IL-2, IL-7, and/or IL-4. (B) Expansion of CD4+ iNKT cells with IL-15 at 5 or 10 ng/mL, with or without cytokine supplementation on day 7 (n=16). Data are shown as mean + SD, and comparisons were analyzed by two-way ANOVA. (C) iNKT cell numbers at baseline (D0) and after 14 (D14) or 21 (D21) days of expansion, with or without autologous PBMCs and α-GalCer added on D14. (D) Glucose and lactate kinetics during iNKT expansion in G-Rex bioreactors.

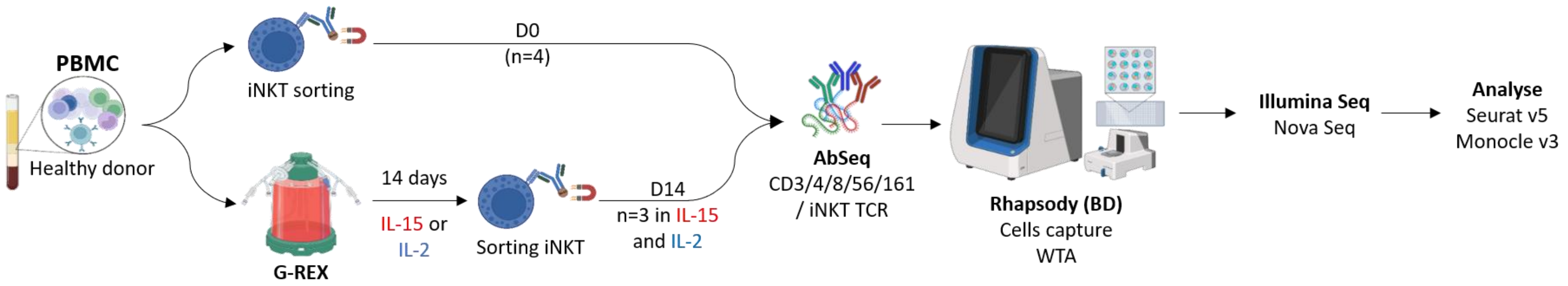


**Supplementary Figure 2: Schematic overview of the single-cell RNA-seq workflow used to analyze human iNKT cells**

iNKT cells (CD3+ iTCR+) were sorted from healthy donor PBMCs either directly ex vivo (D0, n=4) or after 14 days of expansion with IL-15 or IL-2 (D14, n=6). All samples were labelled with AbSeq, captured on the BD Rhapsody platform, subjected to Illumina whole-transcriptome sequencing, and analyzed using downstream mentioned bioinformatic pipelines.

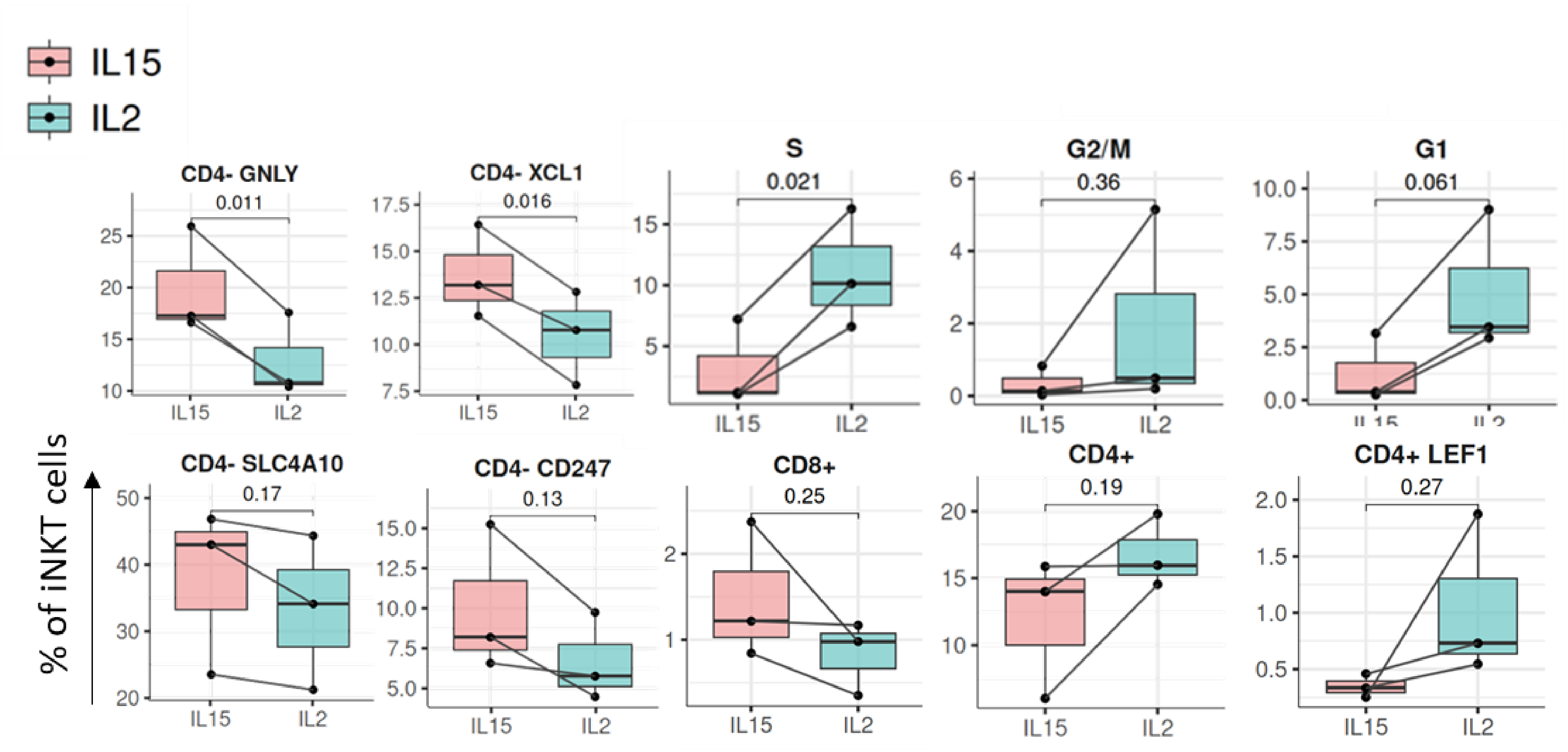


**Supplementary Figure 3: Distribution of iNKT cell clusters in IL-15 versus IL-2 conditions.**

Proportions of iNKT cells in each cluster under IL-15 (red) and IL-2 (blue) conditions. Statistical significance was assessed using a paired t-test.

**A**

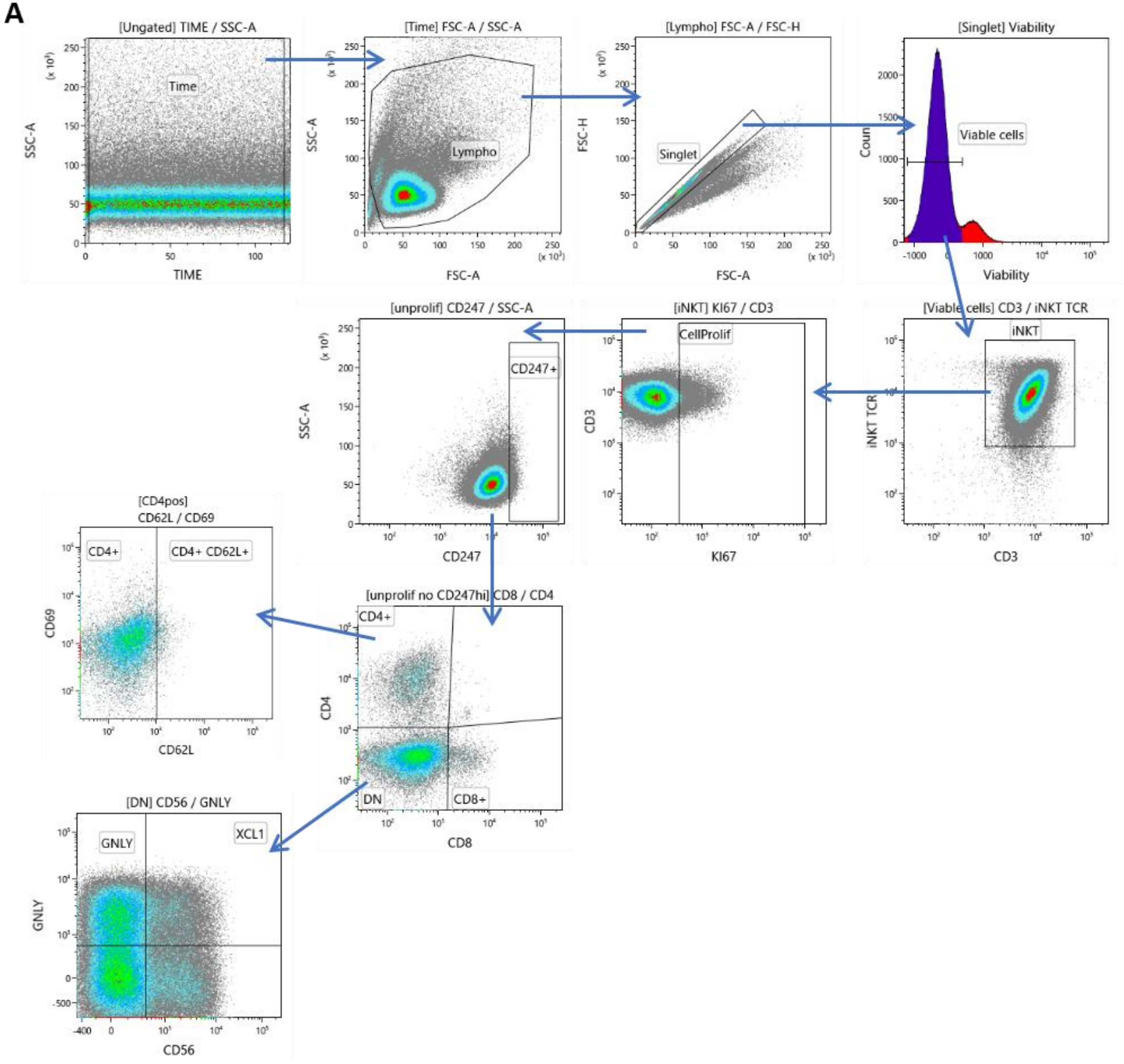


**B**

| Clusters names | Flow cytometry phenotypes |
|---|---|
| CellCycle | iNKT / Ki67+ |
| CD247 | iNKT / Ki67- / CD247high |
| CD8+ | iNKT / Ki67- / CD247low/ CD8+/CD4- |
| CD4+ | iNKT / Ki67- / CD247low/ CD8-/CD4+/CD62L- |
| CD4+ LEF1 | iNKT / Ki67- / CD247low/ CD8-/CD4+/CD62L+ |
| CD4- GNLY | iNKT / Ki67- / CD247low / CD8-/ CD4-/GNLY+/CD56- |
| CD4- XCL1 | iNKT / Ki67- / CD247low/ CD8-/ CD4-/GNLY+ / CD56+ |
| CD4- | iNKT / Ki67- / CD247low/ CD8-/ CD4-/GNLY- |

**Supplementary Figure 4: Gating strategy for the identification of transcriptomic-defined iNKT clusters within flow cytometry data.**

(A) Gating strategy used to identify all iNKT cell clusters after expansion. (B) Summary table outlining the phenotype associated with each cluster.

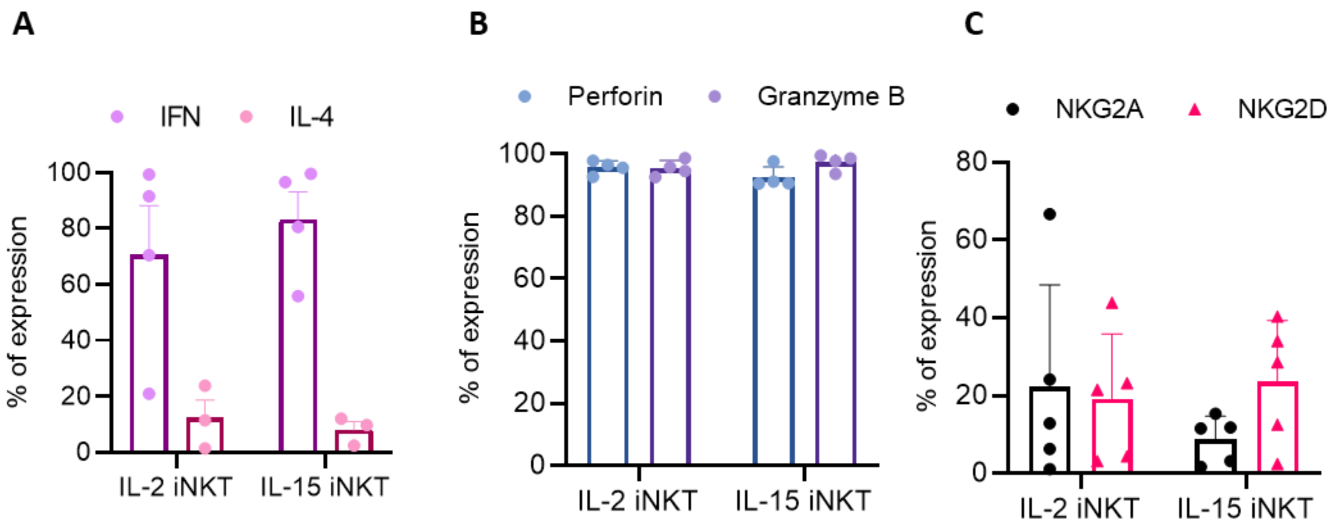


**Supplementary Figure 5: IL-15-expanded iNKT cells exhibit comparable functional potency to IL-2-expanded cells.**

(A) Cytokine expression in total iNKT cells expanded with IL-2 or IL-15 after 4 h of PMA/ionomycin stimulation: IFNγ (n=4) and IL-4 (n=3). Results are shown as mean + SEM. (B) Perforin and granzyme B expression in total iNKT cells after expansion with IL-2 or IL-15 (n=4), presented as mean + SD. (C) NKG2A and NKG2D expression in total iNKT cells after expansion with IL-2 or IL-15 (n=4), presented as mean + SD.

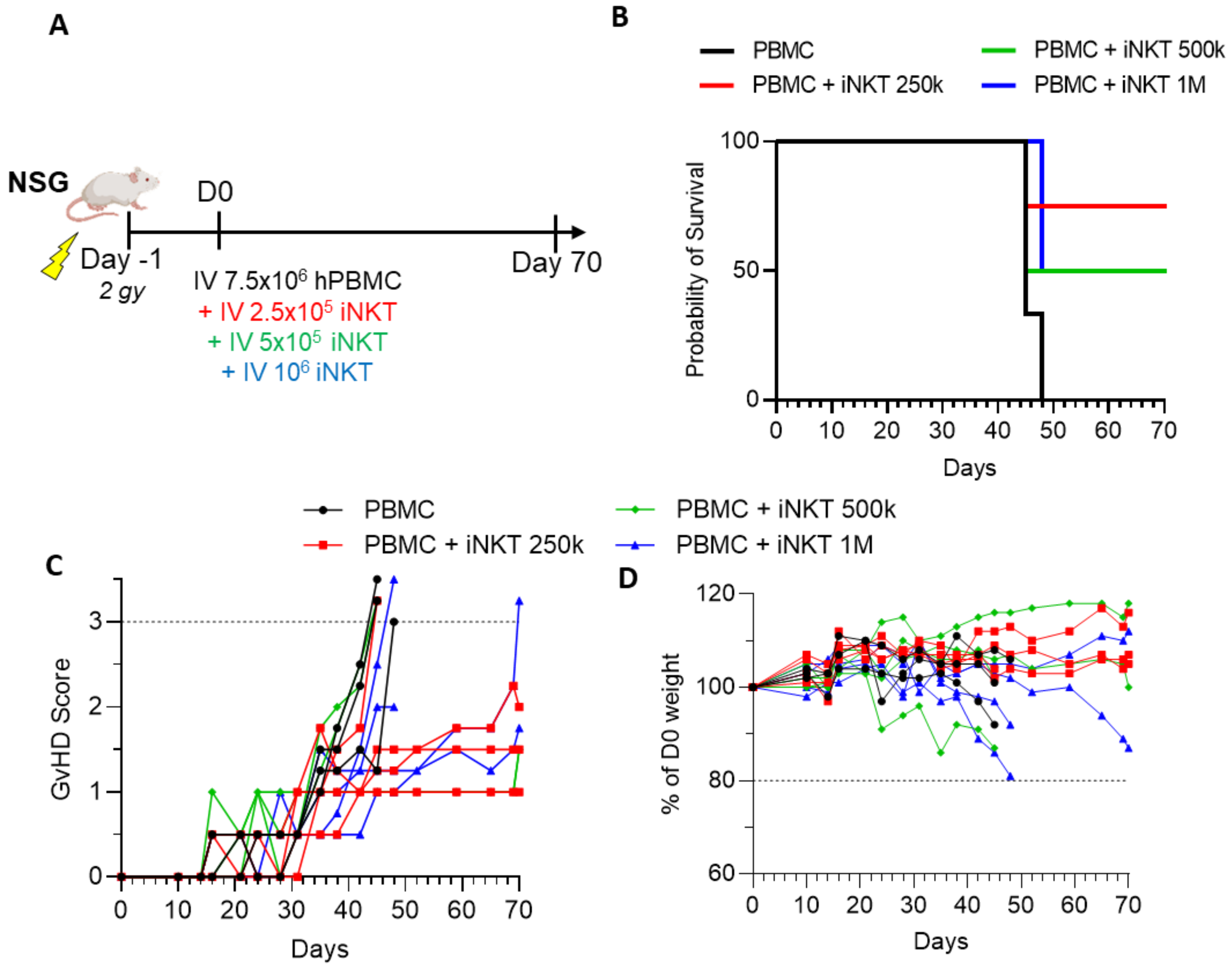


**Supplementary Figure 6: Dose-response analysis of iNKT cells in the GvHD model**

(A) Schema of in vivo study evaluating the anti-GvHD activity of different iNKT cell doses in NSG mice. Mice received 2 Gy irradiation 24 h before injection of $7.5 \times 10^6$ human PBMCs alone (n=3) or together with $1 \times 10^6$ (n=4), $5 \times 10^5$ (n=4), or $2.5 \times 10^5$ (n=4) expanded human iNKT cells. (B) Kaplan-Meier survival of NSG mice after GvHD induction with PBMCs alone (n=3) or PBMCs plus $1 \times 10^6$ (n=4), $5 \times 10^5$ (n=4), or $2.5 \times 10^5$ (n=4) expanded iNKT cells. (C) GvHD clinical scores and (D) body weight, expressed as a percentage of D0, were monitored for 70 days in the PBMC-only group and in PBMC groups receiving the indicated doses of expanded iNKT cells.

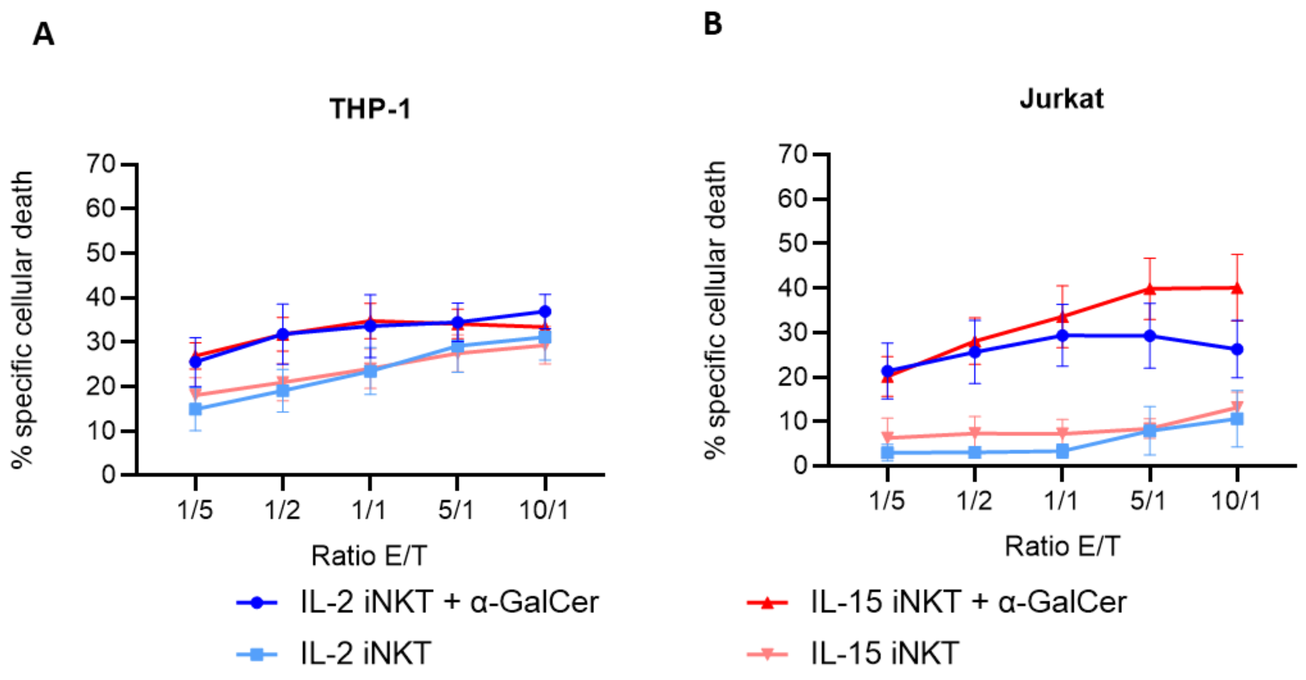


**Supplementary Figure 7: Comparable anti-leukemia cytotoxic activity in vitro between total IL-15 and IL-2-expanded iNKT cells.**

In vitro cytotoxicity of iNKT cells expanded with IL-2 or IL-15 against (A) THP-1 and (B) Jurkat cells (n=8), assessed in the presence or absence of α-GalCer. Results are presented as mean ± SEM.

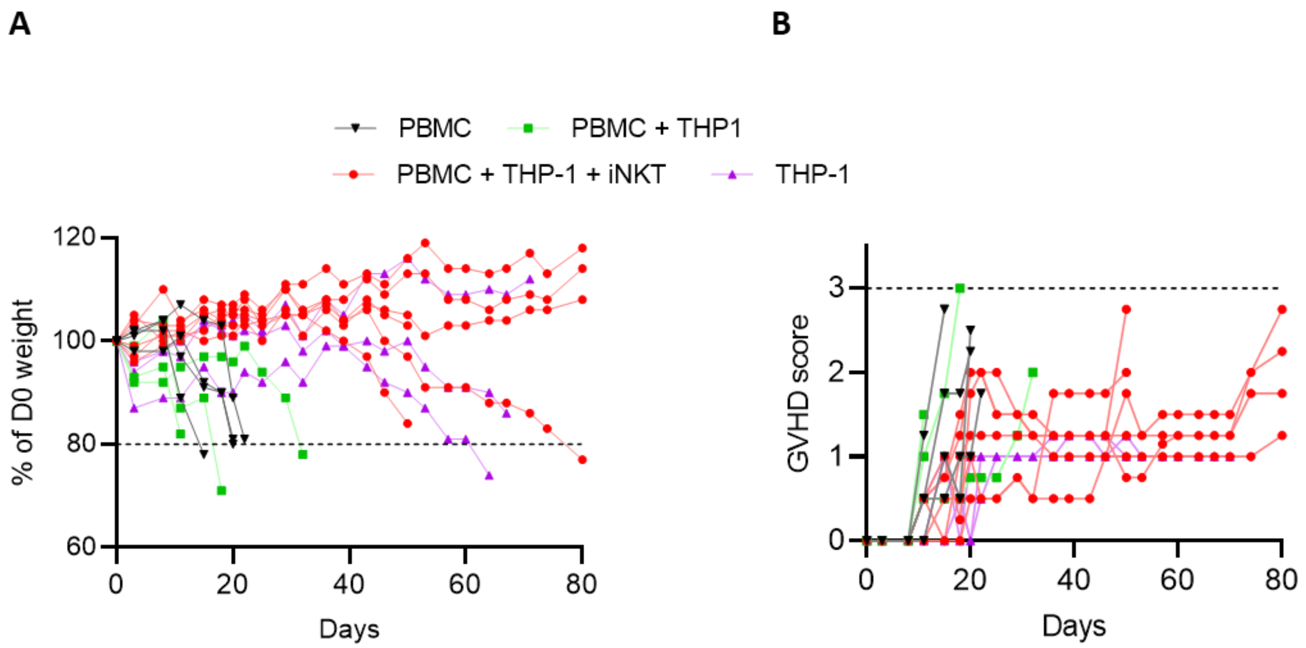


**Supplementary Figure 8: GVHD clinical evolution in the in vivo GVH/GVL model**

Changes in (A) body weight, expressed as a percentage of D0, and (B) GvHD clinical scores across mice treatment groups over the 80 days following transplantation.
